# Anoxia activates CRISPR-Cas immunity in the intestine

**DOI:** 10.1101/2025.04.30.651551

**Authors:** Ian W. Campbell, David W. Basta, Franz G. Zingl, Emily J. Sullivan, Sudhir Doranga, Matthew K. Waldor

**Affiliations:** Division of Infectious Diseases, Brigham and Women’s Hospital, Boston, MA, USA; Department of Microbiology, Harvard Medical School, Boston, MA, USA; Department of Pathology, Brigham and Women’s Hospital, Harvard Medical School, Boston, MA, USA; Howard Hughes Medical Institute, Boston, MA, USA

## Abstract

The natural context in which CRISPR-Cas systems are active in Enterobacteriaceae has remained enigmatic. Here, we find that the *Citrobacter rodentium* Type I-E CRISPR-Cas system is activated by the oxygen-responsive transcriptional regulator Fnr in the anoxic mouse intestine. Since Fnr-dependent regulation is predicted in ~41% of Enterobacteriaceae *cas3* orthologs, we propose that anoxic regulation of CRISPR-Cas immunity is an adaptation that protects Enterobacteriaceae against threats arising from the intestinal microbiome.

## Main

Prokaryotes use CRISPR-Cas systems to recognize and cleave foreign nucleic acid sequences to protect against phages and other mobile genetic elements ^1–3^. However, with a few exceptions ^4,5^, little is known about the regulation of these systems under physiological conditions. The limited knowledge of CRISPR-Cas regulation is partly attributable to the absence of native CRISPR-Cas activity in cultured Enterobacteriaceae, a commonly studied family of bacteria, necessitating investigation using artificial overexpression systems.

*Citrobacter rodentium* is a Gram-negative bacterial pathogen that naturally infects and causes colitis in mice ^6^. Like most Enterobacteriaceae, *C. rodentium* is a facultative anaerobe, capable of growth in the presence or absence of oxygen (oxic or anoxic conditions, respectively). We performed RNA-sequencing of the pathogen’s transcriptional response to anoxia and observed more transcripts from the pathogen’s Type I-E *cas* locus in anoxic vs oxic culture conditions (Fig. 1a; Table S1). To test whether increased *cas* expression correlates with CRISPR-Cas activity we designed a functional assay that monitors the retention frequency of either a plasmid containing a sequence that is (target) or is not (control) recognized by the native *C. rodentium* CRISPR locus (plasmid retention assay; Fig. 1b).

**Fig. 1.**
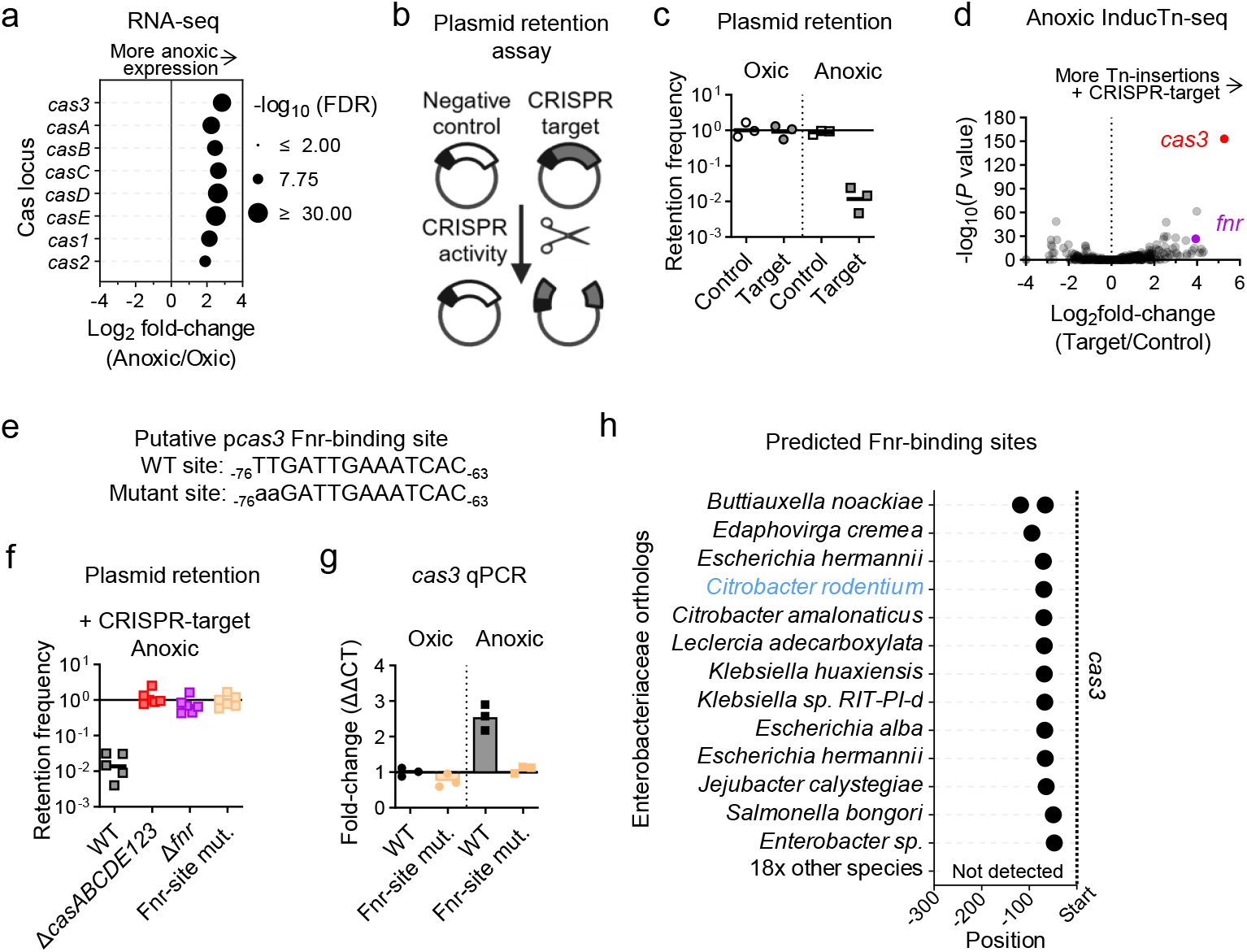
Anoxia causes Fnr-dependent activation of CRISPR-Cas immunity in *C. rodentium*. **a**, RNA-sequencing results from the *cas* locus of *C. rodentium* cultured with (oxic) or without oxygen (anoxic) for 3.5 hours on solid LB agar. The average fold-change of 3 biological replicates is shown. False discovery rate, FDR. **b**, Diagram of plasmid retention assay. Excluding a plasmid with a protospacer target sequence recognized by the native *C. rodentium* CRISPR-Cas system reduces antibiotic-resistant cells. A protospacer adjacent motif (PAM) followed by an unrecognized CRISPR protospacer was used as a negative control. **c, f**, Fraction of cells from a single colony cultured for 24 hours on solid LB agar that retained the target or control plasmid. The geometric mean of biological replicates (colonies; ≥3) are shown. **d**, InducTn-seq volcano plot comparing fold change in the gene insertion frequency between cells that retained the CRISPR target or control plasmid. Points represent individual genes. *P* value from Mann-Whitney U-test. **e**, Sequence of the putative WT and mutated Fnr-binding site relative to the *cas3* start codon. **g**, Quantitative PCR of *cas3* following 3.5 hours of culture on solid LB agar. ΔΔCT (threshold cycle) analysis compared to *rpoA* and oxic culture. 3 biological replicates (plates) and 3 technical replicates per sample. **h**, The location of putative Fnr binding motifs relative to the translational start codon of *cas3* in Enterobacteriaceae orthologs.

In oxic culture conditions, 92% of cells retained the target plasmid, indicating that CRISPR-Cas immunity is inactive (Fig. 1c). In contrast, only 1% of cells retained the target plasmid during anoxic culture (Fig. 1c), demonstrating anoxic-specific immunity. To verify that this assay requires CRISPR-Cas activity, we deleted the entire *C. rodentium cas* locus (Δ*casABCDE123*) and repeated the assay. Deletion of *casABCDE123* eliminated CRISPR-Cas immunity during anoxic culture (Fig. 1f). We conclude that anoxia is required for both expression and activity of *C. rodentium* CRISPR-Cas immunity.

We leveraged the plasmid retention assay for a transposon-insertion loss-of-function screen (InducTn-seq ^7^) to discover anoxic regulators of the *C. rodentium cas* locus. We anoxically cultured a transposon mutant population of *C. rodentium* carrying the target plasmid and then sequenced the mutants that retained the plasmid following antibiotic selection. Compared to a population containing a control plasmid, genes that positively or negatively mediate anoxic CRISPR-Cas immunity are either enriched or depleted in the mutant population containing the target plasmid, respectively (Fig. 1d; Table S2). As expected, transposon insertions in the endonuclease *cas3* were specifically enriched in the population retaining the target plasmid, indicating that disruption of *cas3* prevents CRISPR-Cas immunity. One of the two most enriched non-*cas*-related genes was *fnr*, which encodes an oxygen-responsive transcriptional regulator widely conserved among facultative anaerobic bacteria ^8^ (Fig. 1d). Fnr activity requires iron-sulfur clusters, and the other most enriched gene was a chaperone involved in the maturation of iron-sulfur cluster-containing proteins (*hscA*), previously demonstrated to be needed for full Fnr activity ^9^ (Fig. 1d). Based on these data, we hypothesized that Fnr regulates CRISPR-Cas expression.

In support of this hypothesis, CRISPR-Cas immunity was eliminated in the absence of *fnr* (Δ*fnr*; Fig. 1f). Mutation of a putative Fnr binding motif centered 69.5 nucleotides upstream of *cas3* (Fig. 1e) also eliminated CRISPR-Cas immunity (Fig. 1f). Furthermore, quantitative PCR (qPCR) demonstrated that mutation of the Fnr-binding site upstream of *cas3* eliminated the transcriptional response of *cas3* to anoxia (Fig. 1g). These results indicate that Fnr directly activates CRISPR immunity during anoxia.

To determine the conservation of Fnr-mediated regulation of CRISPR-Cas immunity, we used OrthoDB ^10^ to select 501 non-redundant Gammaproteobacteria genomes containing *cas3* orthologs and interrogated the 300 nucleotides upstream of *cas3* with motif enrichment analysis ^11^. 141 of 501 genomes contained at least one predicted Fnr binding site upstream of *cas3* (Table S3, S4). These genomes were distributed amongst most orders of Gammaproteobacteria (Fig. S1). Notably, the Fnr binding sites in Enterobacteriaceae were primarily centered at the same position as in *C. rodentium* (Fig. 1h, Fig. S2). The positional conservation of an Fnr binding motif in 13 out of 32 Enterobacteriaceae suggests conserved Fnr-dependent *cas3* activation within a subset of this family.

One common anoxic environment encountered by Enterobacteriaceae is within the mammalian intestine. We hypothesized that activation of CRISPR-Cas immunity by anoxia may protect Enterobacteriaceae from threats encountered within the microbially rich intestine. Consistent with this hypothesis, RNA-sequencing revealed that fecal-associated *C. rodentium* isolated from infected C57BL/6J mice had more transcripts from the *cas* locus than in oxic culture (Fig. 2a).

**Fig. 2.**
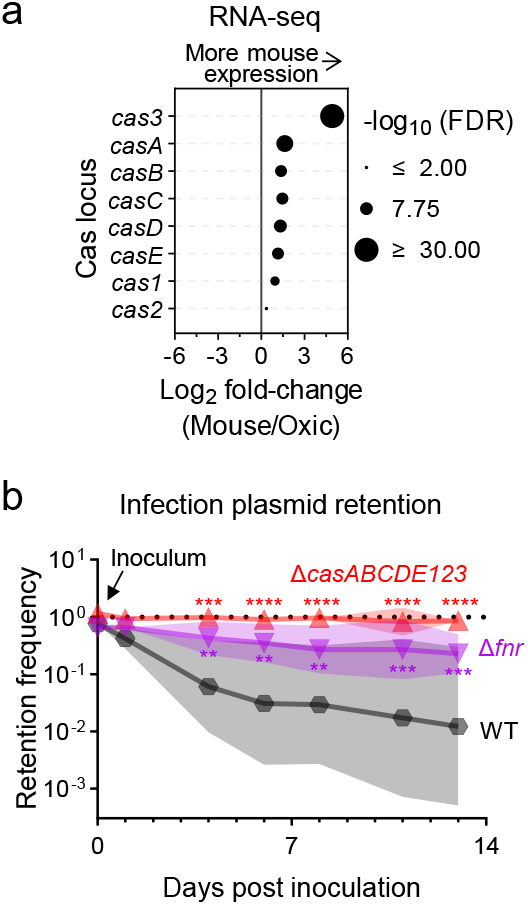
*C. rodentium* CRISPR-Cas immunity is activated by Fnr within the murine intestine. **a**, Cas transcripts from RNA-sequencing of *C. rodentium* recovered from the feces of infected, female, C57BL/6J mice 7 days post inoculation compared to bacteria from oxic culture. 3 animal and 3 oxic samples. **b**, Retention of a CRISPR target plasmid by the indicated strains of *C. rodentium* measured by serial dilution and plating of feces from infected mice (*N* = 16 wild type, 8 Δ*fnr*, 8 Δ*casABCDE123* infected mice; equal mix of sexes). Significance compared to wild type by 2-way ANOVA with Dunnett’s multiple comparison test on log^10^ transformed data.

To determine if this increased *cas* expression results in CRISPR-Cas immunity within the intestine, we infected mice with *C. rodentium* strains carrying the CRISPR-Cas target plasmid and monitored plasmid retention in fecal bacteria. Over the first 24-hours, most shed bacteria retained the plasmid (Fig. 2b). Subsequently, wild-type *C. rodentium* progressively lost the plasmid, with only 1% of shed bacteria retaining the plasmid 13-days post-inoculation. By contrast, 85% of Δ*casABCDE123* cells and 23% of Δ*fnr* cells retained the plasmid over the same 13-day period (Fig. 2b). These results suggest that anoxia within the intestine results in Fnr-dependent activation of CRISPR-Cas immunity in *C. rodentium*. Furthermore, the discrepancy between the Δ*casABCDE123* and Δ*fnr* mutants suggests that there may be intestinal signals in addition to anoxia that regulate CRISPR activity.

Environments like the intestine contain high levels of microbial metabolic activity, which locally depletes oxygen ^12^. Anoxia thus correlates with an increased likelihood of phage and other mobile genetic elements. We therefore propose that anoxic regulation of CRISPR-Cas immunity in *C. rodentium* and other Enterobacteriaceae is an adaptation that protects these bacteria against predation by host-associated threats derived from the microbiome.

## References

1. Sorek, R., Kunin, V. & Hugenholtz, P. Nature Reviews Microbiology 2008 6:3 6, 181–186 (2008).

2. Karginov, F. V. & Hannon, G. J. Mol Cell 37, 7–19 (2010).

3. Marraffini, L. A. & Sontheimer, E. J. Nature Reviews Genetics 2010 11:3 11, 181–190 (2010).

4. Zakrzewska, M. & Burmistrz, M. Front Microbiol 14, 1060337 (2023).

5. Price, V. J. et al. mSphere 4, (2019).

6. Mundy, R., MacDonald, T. T., Dougan, G., Frankel, G. & Wiles, S. Cell Microbiol 7, 1697–1706 (2005).

7. Basta, D. W. et al. Nature Microbiology 2025 1–13 (2025) doi:10.1038/s41564-025-01975-z.

8. Spiro, S. Antonie Van Leeuwenhoek 66, 23–36 (1994).

9. Mettert, E. L., Outten, F. W., Wanta, B. & Kiley, P. J. J Mol Biol 384, 798–811 (2008).

10. Kuznetsov, D. et al. Nucleic Acids Res 51, D445–D451 (2023).

11. Bailey, T. L. & Grant, C. E. bioRxiv 2021.08.23.457422 (2021) doi:10.1101/2021.08.23.457422.

12. Singhal, R. & Shah, Y. M. Journal of Biological Chemistry 295, 10493–10505 (2020).

