## Supplementary material for "Anoxia activates CRISPR-Cas immunity in the intestine": Online methods

### Bacterial Strains

The strains used in this study are listed in Table S5. *Citrobacter rodentium* is a spontaneous streptomycin-resistant isolate of strain ICC168 [1]. *Escherichia coli* strain MFD*pir* was used for cloning [2].

### Oxic and anoxic culture

Bacteria were cultured at 37 °C in either liquid lysogeny broth (LB) shaking at 200 rotations per minute or on solid LB containing 1.5% agar. Oxic culture was performed in atmospheric conditions. Anoxic culture was performed in a Baker Concept 400M anaerobic workstation set to 0% oxygen with media acclimated to the anoxic environment for at least one overnight.

### Plasmid assembly

Plasmid fragments were amplified from plasmid or genomic DNA using single-stranded DNA primers (Integrated DNA Technologies). Fragments were assembled with NEBuilder HiFi DNA Assembly Master Mix (New England Biolabs). Assembled plasmids were transformed into *E. coli* strain MFD*pir* with electroporation and transferred to the recipient strains by conjugation.

### Constructing mutant *C. rodentium* strains

The allelic exchange protocol from Lazarus *et al.,* 2019 [3] was used to create in-frame deletions and the Fnr-site mutation in *C. rodentium*. pTOX5 (Genbank MK972845) was linearized with the restriction enzyme SwaI and assembled with ~1 kb homology arms flanking the desired mutation. Primer sequences are included in Table S6. For deletions, 2-3 codons were left intact at both ends of the deletion. This plasmid was electroporated into MFD*pir*, checked by PCR, and conjugated into *C. rodentium*. Transconjugants were purified by plating consecutively 3-times. Individual colonies were cultured for 1-hour without antibiotic selection and then counter-selected to isolate double crossovers lacking the plasmid backbone. Single colonies were selected, and PCR and whole-genome sequencing confirmed the identity and fidelity of the mutant strains.

### Animal experiments

Animal studies were conducted at Brigham and Women's Hospital in compliance with the 'Guide for the Care and Use of Laboratory Animals' and according to protocols reviewed and approved by the Brigham and Women's Hospital's Institutional Animal Care and Use Committee under protocol 2016N000416. Adult (9–12 weeks) C57BL/6J mice were purchased from Jackson Laboratory (strain #000664) and acclimated for at least 72-hours before experimentation. During infection, mice were housed under specific pathogen-free conditions at 68–75 °F, with 30-50% humidity and a 12-hour light/dark cycle in a biosafety level 2 (BSL2) facility.

For *C. rodentium* infections, mice were deprived of food for 3-5 hours before inoculation. Animals were then mildly sedated with isoflurane and 100 µL of the indicated strain suspended in phosphate-buffered saline (PBS) was inoculated into the stomach with a sterile feeding needle (Cadence Science). Dose (streptomycin-resistant CFU) and CRISPR plasmid retention (gentamicin-resistant CFU) were determined retrospectively by serial dilution and plating.

Animal health was monitored during infection by measuring weight, body condition, and fecal appearance. *C. rodentium* colonization and plasmid retention were monitored by sampling feces from infected animals. Fresh fecal pellets were suspended in sterile PBS and homogenized in a bead beater (BioSpec Products, Inc) with 3.2 mm stainless-steel beads. *C. rodentium* concentration (CFU/g) and plasmid retention were determined by serial dilution and plating for CFUs.

**RNA-sequencing**

For cultured samples, *C. rodentium* was grown overnight in liquid LB in oxic conditions. $4 \times 10^7$ colony forming units (CFU) from the stationary phase culture were seeded onto LB agar plates and cultured for 3.5 hours in the presence or absence of oxygen. Cells were then diluted immediately in 2 parts Qiagen RNAprotect Bacteria reagent and frozen at -80 °C until processing.

For fecal samples, female mice were infected with $5 \times 10^9$ CFU of *C. rodentium* and colonization was monitored in the feces to ensure engraftment. 7-days post-inoculation, fresh feces from infected mice were submerged in RNAprotect Bacteria reagent and manually disrupted with a sterile rod to release bacteria. To remove large debris and eukaryotic cells samples were passed through a 5 μm filter and bacteria were frozen at -80 °C until processing.

RNA was released from bacteria using lysozyme and proteinase K digestion. RNA was extracted with a Qiagen RNeasy kit and purified with RNA Clean & Concentrator-5 (Zymo Research). SeqCenter (Pittsburgh, PA, USA) performed library preparation and sequencing using the following method, provided by SeqCenter: "Samples were DNAse treated with Invitrogen DNAse (RNAse free). Library preparation was performed using Illumina's Stranded Total RNA Prep Ligation with Ribo-Zero Plus kit and 10bp unique dual indices (UDI). Sequencing was done on a NovaSeq X Plus, producing paired end 150bp reads. Demultiplexing, quality control, and adapter trimming was performed with bcl convert".

RNA-sequencing data was processed using CLC Genomics Workbench (Qiagen). RNA-Seq Analysis 2.8 parameters: reference – ICC168 reference genome; mismatch cost = 2; insertion cost = 3; deletion cost = 3; length fraction = 0.8; similarity fraction = 0.8; global alignment; strand specific = reverse; max hits per read = 10; count paired reads as two = no; ignore broken pairs. Differential gene expression was determined with Differential Expression for RNA-Seq 2.8. Results are included in Table S1.

**Plasmid retention assay**

The CRISPR-target plasmid contains the gentamicin resistance gene *aaC1* and a protospacer adjacent motif (PAM) followed by a protospacer sequence matching the native CRISPR array. The control plasmid is identical, except that the protospacer sequence is switched with a protospacer recognized by a different bacterium, and not the native CRISPR-Cas system.

For plasmid retention assays in culture, strains carrying the target or control plasmids were outgrown overnight in an oxic environment on solid LB plates with gentamicin. The next day, strains were restreaked onto LB plates without antibiotics and cultured in the presence or absence of oxygen for 24-hours. Single colonies were resuspended in sterile PBS, and serial dilution was used to determine the fraction of the population that retained the plasmid (gentamicin-resistant divided by streptomycin-resistant CFU).

For plasmid retention assays during infection, an equal mix of male and female mice were inoculated with ~5 x $10^9$ CFU of the indicated strain. Plasmid retention was measured in the inoculum and feces throughout the infection.

**InducTn-seq**

Control or CRISPR-target plasmids were conjugated into a *C. rodentium* InducTn-seq mutant library [4] . 3 x $10^8$ transconjugants were expanded in oxic conditions on LB containing gentamicin, to select for the plasmid, and arabinose, to induce further miniTn5 transposition. The mutant libraries were stored at -80°C in PBS with 20% glycerol. Subsequently, 5 x $10^7$ CFU of the mutant libraries were seeded onto LB agar plates and cultured under anoxic conditions for 24-hours. The population was then expanded in oxic conditions on LB plates containing gentamicin to select for mutant cells that retained the plasmid during anoxic culture. Cells were frozen at -80 °C until processing.

Sequencing libraries were prepared with the protocol from Basta & Campbell *et. al.,* 2025 [4]. Genomic DNA was extracted using a DNeasy Blood and Tissue Kit (Qiagen), sheared to 400 bp using a M220 ultrasonicator (Covaris), repaired using Quick Blunting kit (NEB), polyadenylated using Taq polymerase and dATP, and Illumina P7 adapters were added using T4 ligase (NEB). The end of the mini-Tn5 transposon within the integrated InducTn-seq vector was removed by double restriction enzyme digestion followed by SPRIselect size-selection. Transposon-adjacent sequences were amplified from 800 ng of DNA by PCR using an Illumina i7 index sequence on the reverse primer. Primer dimers were removed by size selection and samples were sequenced on a NextSeq 1000 (Illumina).

InducTn-seq data were analyzed with the protocol from Basta *et. al.,* 2025 [4] using Python. MiniTn5 transposon-insertion frequency was compared between populations that retained the control or target plasmids. Significance was measured with the non-parametric Mann-Whitney U statistical test with Benjamini-Hochberg multiple testing correction. Results are included in Table S2.

**Quantitative PCR**

Bacteria were cultured in liquid LB in oxic conditions. ~$10^7$ colony forming units (CFU) from the culture were seeded onto LB agar plates and cultured for 3.5 hours in the presence or absence of oxygen. After culture, cells were diluted immediately in 2 parts Qiagen RNAprotect Bacteria reagent and frozen at -80 °C until processing.

RNA was released from bacteria using lysozyme and proteinase K digestion, and extracted using an RNeasy kit (Qiagen). qPCR was performed using the Luna Universal One-Step RT-qPCR kit on a StepOnePlus Real-

Time PCR System (Applied Biosystems). At least 3 biological and 3 technical replicates were included per sample, with primers targeting both *rpoA* and *cas3* transcripts. Primer sequences are included in Table S6.

qPCR data were analyzed by comparative critical threshold (CT) analysis. The average *cas3* CT of 3 technical replicates was first normalized to the CT of *rpoA* from the same sample. Then, the CT was normalized using the *cas3* CT from oxic culture, producing ΔΔCT.

**Phylogenetic analysis of *cas3* orthologs**

578 *cas3* orthologs from 500 Gammaproteobacteria were selected for analysis by OrthoDB (version 12.0) [5]. We added *E. coli* strain EDL933 to this list. Strains are listed in Table S3. To construct a phylogeny, we retrieved the nucleotide sequence of *dnaA* from NCBI for 482 of the Gammaproteobacteria and used iTOL (Interactive Tree of Life; version 7.1) [6].

For motif enrichment analysis, we retrieved 300 bp upstream of *cas3* from NCBI for 553 of the 579 *cas3* orthologs. These sequences were compared to the prokaryotic transcription factor motif database PRODORIC (release 2021.9) with Simple Enrichment Analysis (SEA; version 5.5.7) [7] with the parameters: differential enrichment analysis; both strands; Fisher Exact Test; control sequences from shuffled sequences, preserving 3-mer frequencies; hold-out 10% of sequences. This analysis determined that the Fnr motif defined in *E. coli* strain MG1655 (ID MX000004) was significantly enriched within the dataset (*P* value 6.87e-5). The location relative to *cas3* and the scores of putative Fnr binding sites are included in Table S4.

**Software and statistics**

Data analysis was performed using CLC genomics workbench (version 24.0.1), GraphPad Prism (version 10.4.1), Python (version 3.12), and Microsoft Excel. The number of samples and statistical tests are described in the figure legends. Graphics were prepared with GraphPad Prism and Microsoft PowerPoint. RNA-seq and InducTn-seq sequencing reads are deposited in the Sequencing Read Archive (SRA) under accession no. PRJNA1254768. Reads were mapped to the *C. rodentium* ICC168 genome FN543502.1.

**Method references**


1. Campbell, I. W., Hullahalli, K., Turner, J. R. & Waldor, M. K. Quantitative dose-response analysis untangles host bottlenecks to enteric infection. *Nature Communications 2023 14:1* **14**, 1–13 (2023).
2. Ferrières, L. *et al.* Silent mischief: Bacteriophage Mu insertions contaminate products of Escherichia coli random mutagenesis performed using suicidal transposon delivery plasmids mobilized by broad-host-range RP4 conjugative machinery. *J Bacteriol* **192**, 6418–6427 (2010).
3. Lazarus, J. E. *et al.* A new suite of allelic-exchange vectors for the scarless modification of proteobacterial genomes. *Appl Environ Microbiol* **85**, (2019).

4. Basta, D. W. *et al.* Inducible transposon mutagenesis identifies bacterial fitness determinants during infection in mice. *Nature Microbiology 2025* 1–13 (2025) doi:10.1038/s41564-025-01975-z.

5. Kuznetsov, D. *et al.* OrthoDB v11: annotation of orthologs in the widest sampling of organismal diversity. *Nucleic Acids Res* **51**, D445–D451 (2023).

6. Letunic, I. & Bork, P. Interactive Tree of Life (iTOL) v6: recent updates to the phylogenetic tree display and annotation tool. *Nucleic Acids Res* **52**, W78–W82 (2024).

7. Bailey, T. L. & Grant, C. E. SEA: Simple Enrichment Analysis of motifs. *bioRxiv* 2021.08.23.457422 (2021) doi:10.1101/2021.08.23.457422.

**Competing interests**

The authors declare no competing interests.

**License Information**

This article is subject to HHMI's Open Access to Publications policy. HHMI lab heads have previously granted a nonexclusive CC BY 4.0 license to the public and a sublicensable license to HHMI in their research articles. Pursuant to those licenses, the author-accepted manuscript of this article can be made freely available under a CC BY 4.0 license immediately upon publication.

**Funding**

Howard Hughes Medical Institute (MKW)
Life Science Research Foundation: Zingl-2024HHMI (FGZ)
NIH grants: P30 DK034854 (IWC), R01 AI042347 (MKW)

**Acknowledgements**

We thank members of the Waldor lab for helpful discussions and feedback on the manuscript.
