## Supplemental figures for "Anoxia activates CRISPR-Cas immunity in the intestine"

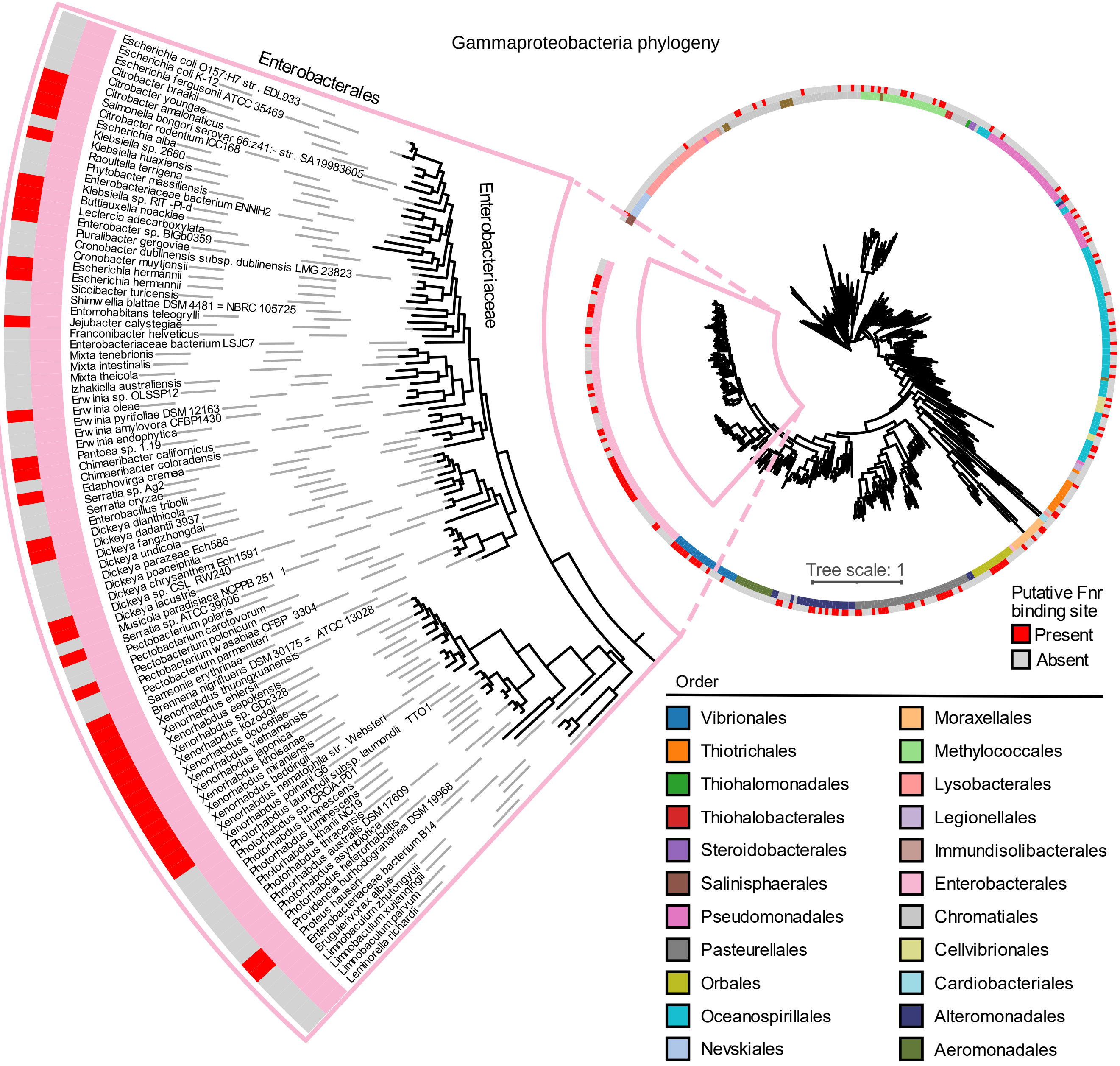


**Supplemental fig. 1 | Potential Fnr binding sites are present upstream of *cas3* in many Gammaproteobacteria genomes.** Gammaproteobacteria *cas3* orthologs were curated from OrthoDB and the phylogeny was constructed from 482 *dnaA* sequences. Putative Fnr binding sites in the 300 bp upstream of *cas3* were identified with motif enrichment analysis with the Fnr motif defined in *E. coli* strain MG1655. Tree scale is the number of substitutions per site. Strains are listed in Table S3 and putative Fnr binding sites are listed in Table S4.

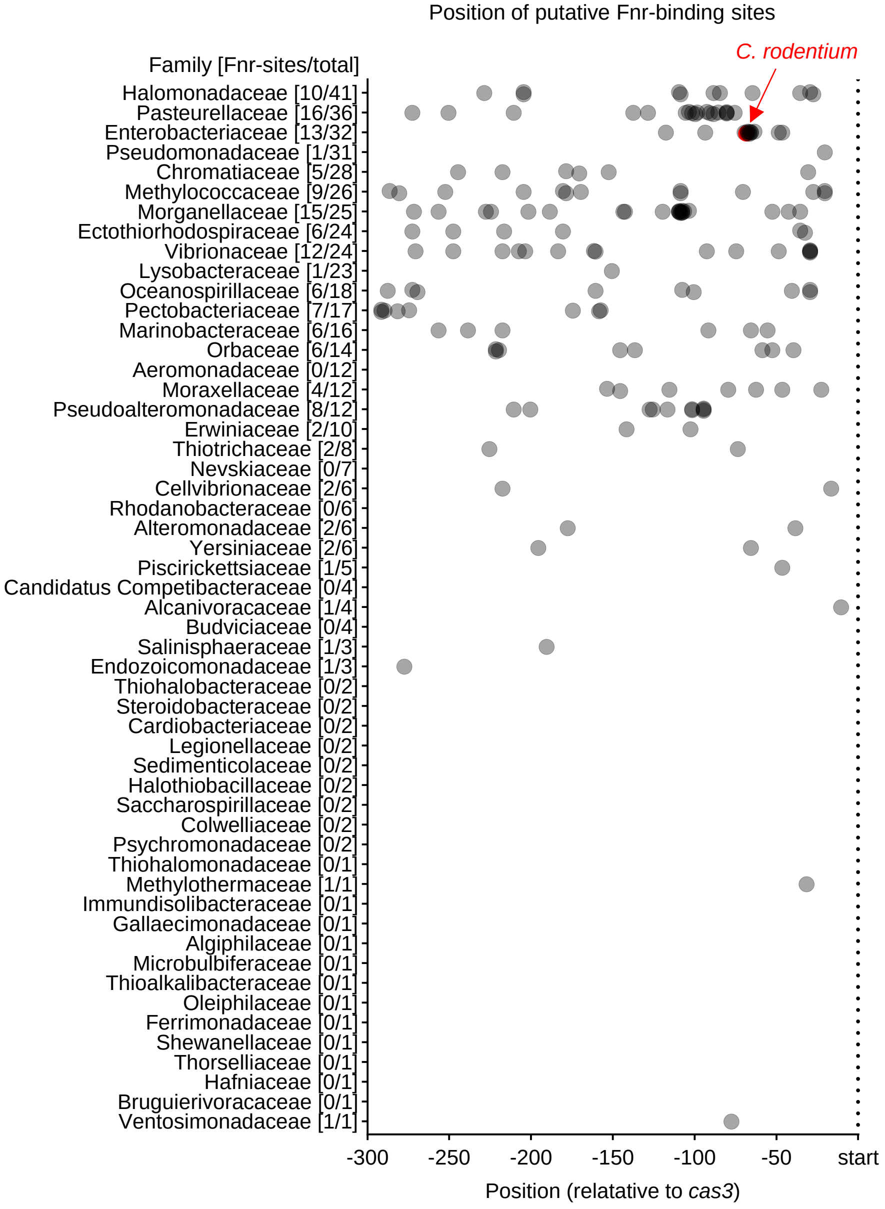


**Supplemental fig. 2 | There is positional conservation of a putative Fnr binding site upstream of *cas3* in 13 of 32 Enterobacteriaceae.** The position of putative Fnr binding sites relative to 553 Gammaproteobacteria *cas3* orthologs, grouped by family. The number of genomes with at least one putative Fnr binding site in the 300 bp upstream of *cas3* (numerator) and the number of genomes analyzed within the family (denominator) are next to the family's name. Strains are listed in Table S3 and putative Fnr binding sites are listed in Table S4.

2
